# A Duplicate Resolved Paddlefish Genome Provides Insights into the Mechanisms of Rediploidisation and Hox Cluster Evolution

**DOI:** 10.64898/2026.08.13.744671

**Authors:** Dearbhaile Casey, Lukasz Niezabitowski, Manu Kumar Gundappa, Hanover Matz, Arun Venugopalan, Larry A. Hanson, Helen Dooley, Daniel J. Macqueen, Anthony K. Redmond, Aoife McLysaght

**Affiliations:** Smurfit Institute of Genetics, Trinity College Dublin, Dublin, Ireland; Animal Breeding and Genomics, Wageningen University & Research, Wageningen, The Netherlands; Department of Microbiology and Immunology, University of Maryland School of Medicine, Institute of Marine and Environmental Technology, Baltimore, MD, USA; Department of Pathobiology and Population Medicine, College of Veterinary Medicine, Mississippi State University, Starkville, MS, USA; Department of Comparative Biomedical Sciences, College of Veterinary Medicine, Mississippi State University, Starkville, MS, USA; The Roslin Institute and Royal (Dick) School of Veterinary Studies, University of Edinburgh, Edinburgh, Scotland; School of Medicine, University College Dublin, Dublin, Ireland

## Abstract

Whole-genome duplication (WGD; or polyploidy) has played a major role in the evolution of many lineages however, our understanding of the processes that shape genome evolution following WGD remains incomplete. While polyploidy duplicates the entire genome sequence, it is rediploidisation that establishes independent duplicated genes. Rediploidisation proceeds through suppression of meiotic recombination across polysomic loci thus restoring disomic inheritance, a process that is not synchronised across the genome. Despite its importance, the mechanisms underlying this process remain poorly understood. The slowly evolving genomes of paleopolyploid Acipenseriformes paddlefish and sturgeon provide an invaluable system for investigating this, as rediploidisation was highly asynchronous in these lineages. In both genomes ohnologs tend to segregate into blocks on the chromosomes according to rediploidisation timing, a pattern that suggests links between chromosomal structure and rediploidisation. Here, we analyse a newly-produced duplicate-resolved paddlefish genome assembly and show a strong concordance between genome rearrangement and rediploidisation timing. We also find that topologically associated domain (TAD) boundaries are associated with rediploidisation block boundaries. Together these results indicate that rediploidisation in acipenseriformes occurred through a process of genome rearrangements that was subject to functional constraints imposed by 3D genome architecture. We investigate the evolution of Hox clusters in these lineages, revealing a previously overlooked duplicate HoxC region in paddlefish, and both ancestral and lineage-specific Hox cluster rediploidisation with substantially different timings. These findings highlight a complex evolutionary history following WGD in Acipenseriformes with implications for understanding short-term adaptations to polyploidy as well as longer-term diversification of lineages.

## Introduction

Ancient whole genome duplications (WGD) have occurred in many eukaroyotic lineages (1, 16, 37, 49, 50, 74, 81, 87, 88). These doubling events are thought to offer raw genetic material with the potential to facilitate evolutionary innovations in descendant species (83). The process of rediploidisation is crucial to unlocking this potential.

WGD can occur by self-doubling of the genome (autopolyploidistaion) or through hybridisation of two different parent species (allopolyploidisation). Cytogenetically, in many cases, the latter may be diploid-on-arrival (i.e. no classical rediploidisation step is required) as the chromosomes are likely to be sufficiently diverged to form preferential bivalent pairs at meiosis. However, in autoploylploids there is no sequence differentiation and the chromosomes will experience polysomic inheritance due to the occurrence of meiotic recombination either multivalently, or in non-preferential bivalents (such that any of the four chromosomes may recombine with any of the others). This situation persists until recombination becomes suppressed. The suppression of recombination establishes independent duplicated loci from the previously polysomic locus; that is, it creates duplicated genes (ohnologs) where previously there were merely additional alleles (70) (Redmond et al. in review) (56). Sxuppression of meiotic recombination is the crux of rediploidisation, establisheing independent duplicated loci, free to undergo uninterrupted sequence and functional divergence (70, 71).

Seen this way, rediploidisation is key to unlocking much of the potential of WGD for evolutionary novelty and diversification (14, 71). Despite being such a critical process in post-WGD eukaryotic evolution, the mechanisms of rediploidisation are poorly understood. The most popular hypothesis is that rediploidisation is brought about by genomic rearrangements and the accumulation of mutations along homologous chromosomes (25, 50). However, evidence in support of genome rearrangements as a mechanism of rediploidisation is scant, though a recent example implicates chromosomal fusions as instigators of rediploidisation in autotetraploid snow carp (90).

Investigating the patterns and mechanisms of rediploidisation requires suitable study systems where the WGD is sufficiently old enough so that rediploidisation has progressed but recent enough so that events can still be tracked through evolutionary time. The shared WGD at the base of the extant Acipenseriformes (sturgeons and paddlefish) provides an ideal system. Noted for their slow evolutionary rate (10, 29, 43) they have recently been established as a model of slow, asynchronous rediploidisation. Despite having shared an autopolyploidisation event well over 200Ma (million years ago), more than half of the genome remained tetraploid (i.e. under tetrasomic inhertiance) until after the divergence of the two acipenseriform lineages (*∼* 170 Ma in the Jurassic) (70). Their genomes are thus a mosaic of ohnolog pairs that rediploidised either before (PreSpec; pre-speciation) or after (PostSpec; post-speciation) the speciation event separating sturgeons and paddlefish known as AORe (Ancestral Ohnolog Resolution) and LORe (Lineage-specific Ohnolog resolution) ohnolog pairs respectively (70, 71). Importantly, ohnolog pairs with a shared rediploidisation history are not randomly distributed across the genome, instead, they form distinct syntenic blocks along uninterrupted sections of chromosomes (70). Based on this, Redmond et al. (2023) proposed a model for rediploidisation where the structural differences introduced by genome rearrangements interrupt meoitic pairing and homologous recombination in that genomic region, such that rediploidisation proceeds rearrangement by rearrangement along and between chromosomes (Fig 1 (a)). This model is conceptually similar to the establishment of mammalian sex chromosomes which formed by punctuated rearrangement events, each progressively suppressing X-Y recombination and establishing ‘evolutionary strata’ of chromosome divergence (31, 44). While rearrangements appear to be a plausible explanation for the stratified topologies of paddlefish and sturgeon given that blocks with shared histories are apparent in visual inspection in a circos plot (1(b)(i)) a formal investigation of this model of rediploidisation is still lacking.

**Figure 1.**
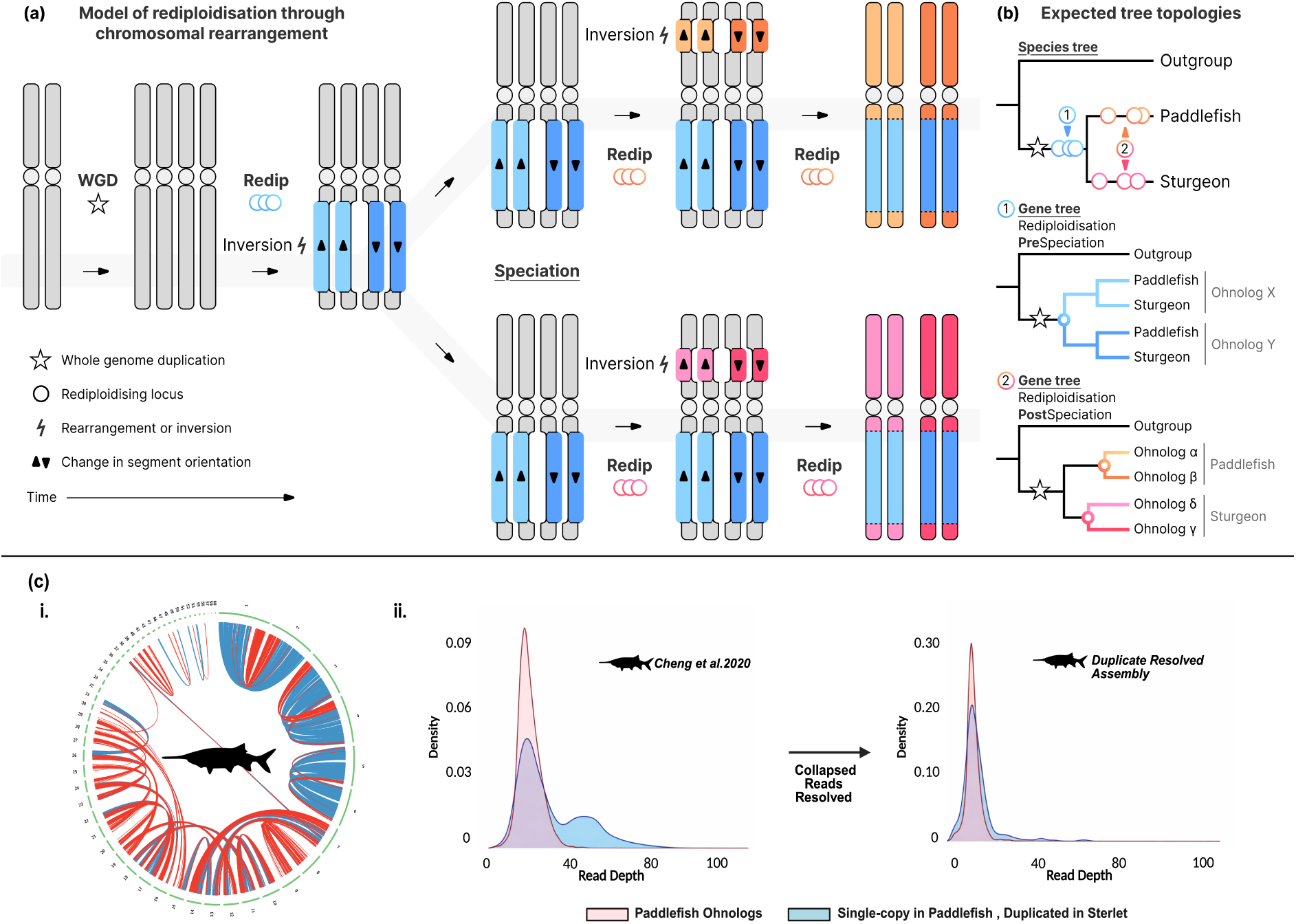
Model for rediploidisation through chromosomal rearrangement and resolving collapsed reads in the paddlefish genome. **(a)** The schematic shows diploid chromosomes that duplicate via WGD, resulting in a tetraploid. A meiotic rearrangement (inversion) suppresses recombination and promotes their rediploidisation (blue loci). These ancestral duplication events give rise to *PreSpec* gene tree topologies in phylogenetic analyses (far right, (b)(1)). Following speciation, independent rearrangements occur in each species, shown in shades of orange for paddlefish and shades of pink for sturgeon. These generate lineage-specific ohnologs and *PostSpec* gene tree topologies (far right, (b)(2)). **(c)(i)** Circos plot showing *PreSpec* ohnolog pairs as blue links and *PostSpec* ohnolog pairs as red links in the Paddelfish Genome (ii) Read-depth distributions for paddlefish ohnolog pairs (red) and putative ohnologs (blue) In the published genome assembly (left), genes present as single copies in paddlefish but duplicated in sterlet show a peak at approximately *∼*2*×* the average coverage, indicating that duplicated loci were collapsed into a single genomic region during assembly. In the duplicate-resolved assembly (right), this peak is substantially reduced. Read depth is expressed as fold coverage (*×*), where 1*×* represents the genome-wide average sequencing coverage per base.

Intriguingly, structuro-functional constraints imposed by 3D genome architecture have been proposed to influence chromosomal rearrangement possibilities, though this has never been linked to the pattern of asynchronous rediploidisation across the genome. For example, topologically associating domains (TADs) are regions of elevated intra-domain chromatin contact that often contain genes with shared transcriptional programmes and are thus thought to play important roles in genome organisation and function (52) Importantly, disruption of TAD boundaries has in some cases been linked to negative outcomes including oncogenesis, embryonic lethality, and developmental disease (58, 82) Some TADs have been demonstrated to be evolutionarily conserved, and to often harbour age-matched genes, including those retained following WGD, and have vitally been shown to be resistant to disruption by genomic rearrangements (38, 64, 84). This raises the possibility that preserving normal 3D genome organisation may selectively constrain viable chromosomal rearrangement space and, consequently, influence the patterns and rates of rediploidisation following WGD.

To address key open questions highlighted above, we first improved the assmebly of the American paddlefish (*Polyodon spathula*) genome by resolving mistakenly collapsed regions (17) and re-annotated it using newly generated RNA-seq data. Assembly enhancement was necessary because these genomes are exceptionally slow evolving, causing many ohnologous genes and genomic regions to be erroneously collapsed into single assembly loci. Ohnologs that underwent rediploidisation after the divergence of paddlefish and sturgeons –and therefore exhibit the lowest sequence divergence-were, as expected, the most severely affected (70). Using this enhanced assembly, we identified micro-and macro-synteny blocks of ohnologs as proxies for rearrangements and found that both rearrangement breakpoints and TAD boundaries coincide with changes in rediploidisation history along the chromosome (which we dub ‘topology breakpoints’; Fig 2 (a)). Patterns of 3D genome architecture appear to constrain the genomic locations at which rearrangements occur, and functional analyses further reveal distinct enrichment patterns among gene categories associated with early and late rediploidisation ohnologs. Building on this, we then investigated the rediploidisation patterns of the acipenseriform *Hox* clusters. Previous work has shown that the duplicate *HoxA*, *HoxB*, and *HoxD* gene clusters are largely retained in Acipenseriformes, whereas the duplicate *HoxC* cluster was inferred to be lost in the paddlefish but retained in sturgeon, echoing similar losses observed in some elasmobranch lineages (12, 15, 41). By reconstructing Hox cluster evolution, we find major differences in rediploidisation histories among clusters, and reveal a previously misassembled, degenerate (rather than lost) duplicate *HoxC* region in paddlefish. We speculate that variation in the timing of *Hox* cluster duplication (that is, rediploidisation) may have contributed to phenotypic divergence between these sister lineages, perhaps consistent with hypotheses linking polyploidy and asynchronous rediploidisation to adaptive potential during periods of environmental instability (70).

**Figure 2.**
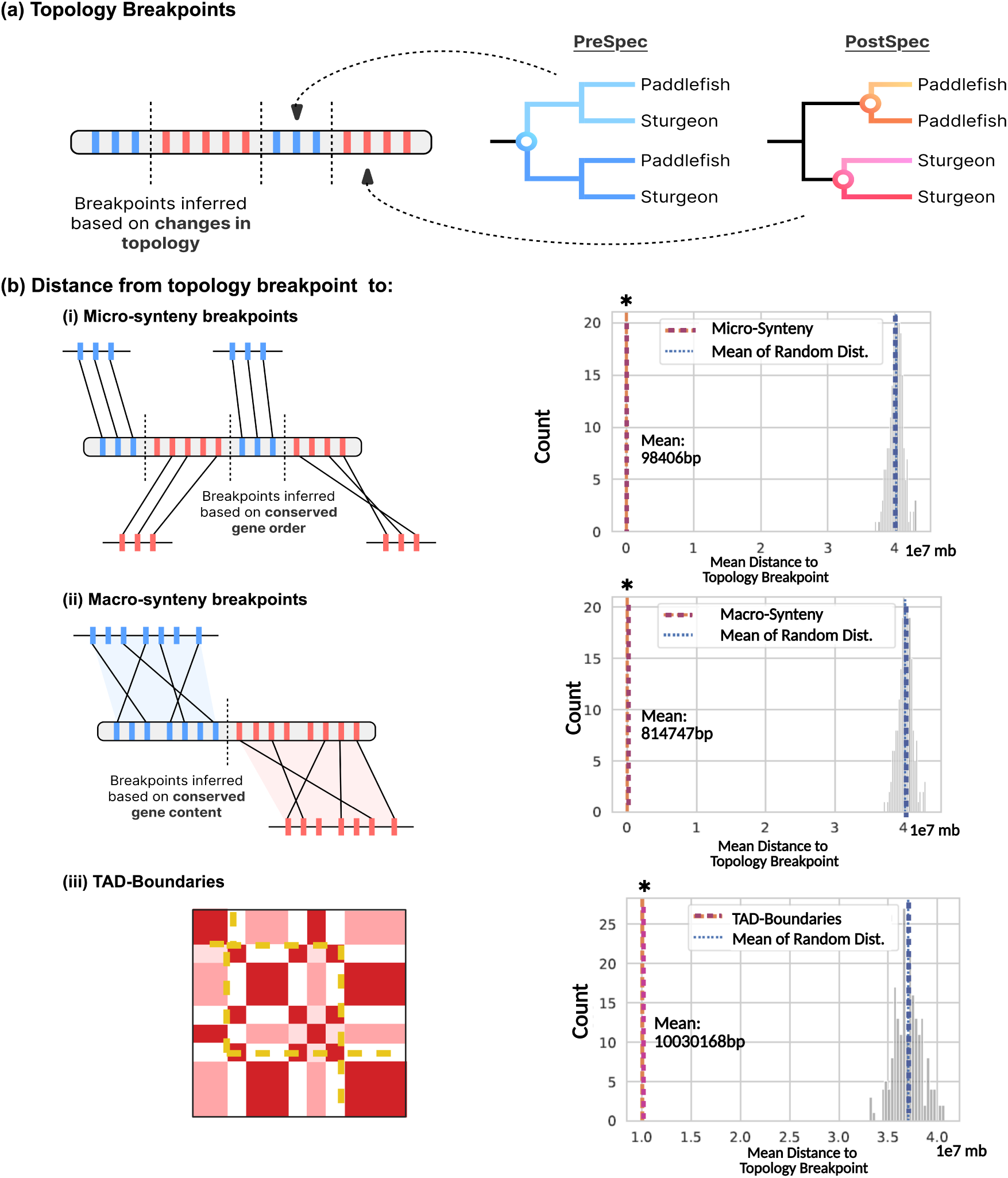
Rearrangements are associated rediploidisation in Acipenseriformes genomes. **(a)** We compared the distance from topology breakpoints to: **(b)(i)** micro synteny blocks, representing smaller and potentially more recent rearrangements; **(ii)** macro-synteny blocks, representing older and larger scale rearrangements and **(iii)** topologically associating domain (TAD) boundaries. Distances from these genomic features to topology breakpoints were compared with distances from randomly sampled genomic locations (only paddlefish shown). Statistical significance was assessed using a one-sided Mann-Whitney U test, and the observed mean distance was evaluated against a null distribution generated from random simulations. Asterix indicates significant differences in mean distance.

## Results

### Characterising ohnologs with a new, duplicate-resolved paddlefish genome

Numerous ohnolog pairs appear to have been erroneously collapsed into into singleton assembly regions in the paddlefish genome (70). In line with this, the sturgeon genome is reported to have retained thousands of genes after WGD that were proposed to be lost in paddlefish, including an additional *HoxC* cluster (6, 15, 17), however, we considered that many of these may in fact be assembly error artegfacts. As such we first sought to produce an improved paddlefish genome assembly in order to determine if the elevated rate of gene loss reported in paddlefish can be wholly or partially ascribed to error.

We found that approximately 2.3% of the paddlefish genome shows elevated read depth, consistent with collapsed regions (Fig 1(c)(ii)). Among genes that are duplicated in sturgeon but recorded as single-copy in paddlefish, there is a secondary read depth peak at double coverage suggesting that the collapsed genomic regions contain ohnolog pairs (Fig 1 (c)(ii); Supplementary Materials (22)). Adapting a reduplication strategy from (17) to avoid ohnologous region collapse we re-assembled the paddlefish genome, and indeed the secondary peak of supposed singletons with double read depth disappears (Fig 1 (c)(iii). This improved assembly and genome annotation was significantly enhanced by supplementing with new RNA-seq data from 15 tissues from 4 individuals and integrating three annotation strategies to predict 35,930 protein-coding genes with BRAKER3 (30) This is a remarkable increase of 9,859 genes compared to the original assembly (12) However, this still falls over 10,000 genes short of the predicted protein-coding gene count in the sterlet sturgeon, implying that although previous estimates of paddlefish protein-coding genes were erroneously low, the gene loss rate after their shared WGD was indeed higher in paddlefish than in sturgeon.

Building on the work of Redmond et al. (70), we classified ohnolog gene trees based on topology as either PreSpec (shared duplication node) or PostSpec (independent duplication nodes). By integrating phylogenetic and syntenic evidence, we recovered patterns consistent with those reported by Redmond et al. (70), while also identifying approximately 423 additional ohnolog pairs. PostSpec topologies were more prevalent than PreSpec topologies, with the remaining gene trees exhibiting intermediate histories (PreSpec-like or PostSpec-like) (Supplementary Table S1), consistent with Redmond et al 2023. The discovery of these additional genes and ohnolog pairs from our new assembly should allow for better analysis of synteny and its link to rediploidisation in paddlefish (70).

### Associating rearrangement breakpoints with rediploidisation event

To identify whether chromosomal rearrangements are associated with rediploidisation events, we first needed to locate syntenic blocks of genes with consistent rediploidisation histories. These blocks are noticeable by eye in the circos plots (Fig 1 (c)(i)), especially on the larger chromosomes, where extended stretches of consecutive blue and red links (joining prespec or postspec ohnolog pairs, respectively) occur sequentially along chromosomes.

To formally identify genomic locations where the syntenic chain of ohnolog pair gene trees changes topology (henceforth referred to as topology breakpoints) we applied a Bayesian segmentation algorithm (Niezabitowski et al., BIORXIV). Briefly, our approach models each chromosome as a sequence of categorical observations (prespec or postspec) with posterior probabilities, and a ‘breakpoint’ is defined as the first element in the sequence where the probability changes between topology category (see NIEZABITOWSKI BIORXIV for a full explanation of this approach). Using this method, we identified a total of 383 topology breakpoints across the 60 largest chromosomes of paddlefish and 293 breakpoints across the 60 largest sterlet chromosomes (Supplementary Table S2; and Figshare (22)).

We used synteny block co-ordinates as a proxy for genome rearrangement breakpoints We identified pairwise micro-synteny blocks between the genomes using i-ADHoRe (67) (Fig 2 (b)(i)). These blocks represent regions where both gene order and orientation have been retained or jointly modified. We also identified macro-synteny blocks shared by both species using a method developed in (61, 62) and adapted in Niezabitowski et al., 2026 (Fig 2 (a)(i&iii)).

We next related these synteny blocks to the topology breakpoints (Fig 2 (b) (ii), (70); Supplementary Materials; Table 1 (22)) to assess whether genome rearrangement events were spatially associated with transitions in rediploidisation history. Specifically we quantified the genomic distance between each synteny block boundary and the nearest topology breakpoint, and compared this to a null distribution generated from random genomic coordinates. This allowed us to test whether shifts between early (PreSpec) and late (PostSpec) rediploidisation topologies along a chromosome, which we take to mark the boundaries of rediploidising units, occur significantly closer to rearrangement boundaries than expected by chance (Fig 2 (a)). We found that our rearrangement blocks were in closer proximity, both in base pair distance and gene distance, to topology breakpoints than to randomly chosen locations in the genomes of both fish (Fig 2 (b) (ii)and(iii), Supplementary Materials (22).)

Finally, in addition to rearrangement block associations we also assessed whether TAD boundaries were associated with topology breakpoints (Fig 2 (b) (iii); Supplementary Materials; Table 1 (22); Supplementary Table S5). Using the logic above, we assessed whether TAD boundaries were associated with topology breakpoints. We find that TAD boundaries are indeed spatially associated with topology breakpoints and the observed proximity of the two therefore suggests that entire chromatin domains, rather than isolated individual genes, tended to rediploidise together, possibly as functional units.

### Functional organisation of early-and late-rediploidising regions

Given this enhanced information on the origin of these rediploidsing units, we examined their chromosomal distribution in greater detail. We classified blocks as PostSpec-type or PreSpec-type topology blocks when they contained a significant enrichment of PostSpec or PreSpec genes relative to the genomic background (one-sided Fisher’s exact test *p <* 0.05; Supplementary Materials; Table 2 S2). The majority of PostSpec-type blocks were located on Paddlefish chromosomes 8, 7, 16, 18 and 22, and Sterlet chromosomes 20 16, 12, 15, and 19. By contrast, PreSpec-type blocks were predominantly found on the 6 largest chromosomes in both species consistent with findings from Redmond et al. (2023) (Supplementary Materials; Table 2 (22)).

We next examined the biological functions enriched within the genes contained in these blocks by performing functional enrichment analyses using STRING on the PreSpec and PostSpec ohnolog sets (78). PreSpec gene sets showed strong and consistent enrichment for pathways associated with chromosome segregation, cell-cycle control, centrosome maturation, homologous recombination, and G2/M checkpoint regulation, precisely the types of functions which may be advantageous in stabilising meiosis, mitosis, and cell biology in the immediate aftermath of WGD (all enriched terms from STRING can be found in the Supplementary Materials (22)). Notably, pathways such as AURKA activation by TPX2, which is a critical allosteric mechanism that drives mitotic spindle assembly centrosome maturation, and chromosome segregation was one of the most significantly enriched. Progesterone mediated oocyte maturation and the MAPK (Mitogen-Activated Protein Kinase) pathway were also enriched. The MAPK pathway is a crucial cell signaling network that relays external signals (like growth factors, stress) from the cell surface to the nucleus, controlling fundamental processes such as cell proliferation, differentiation apoptosis, and stress response. In contrast, PostSpec gene sets showed no clear pattern of enrichment: PPI networks were not significantly enriched and functional terms were varied and showed no clear consistency (Fig3(c) and (d)).

**Figure 3.**
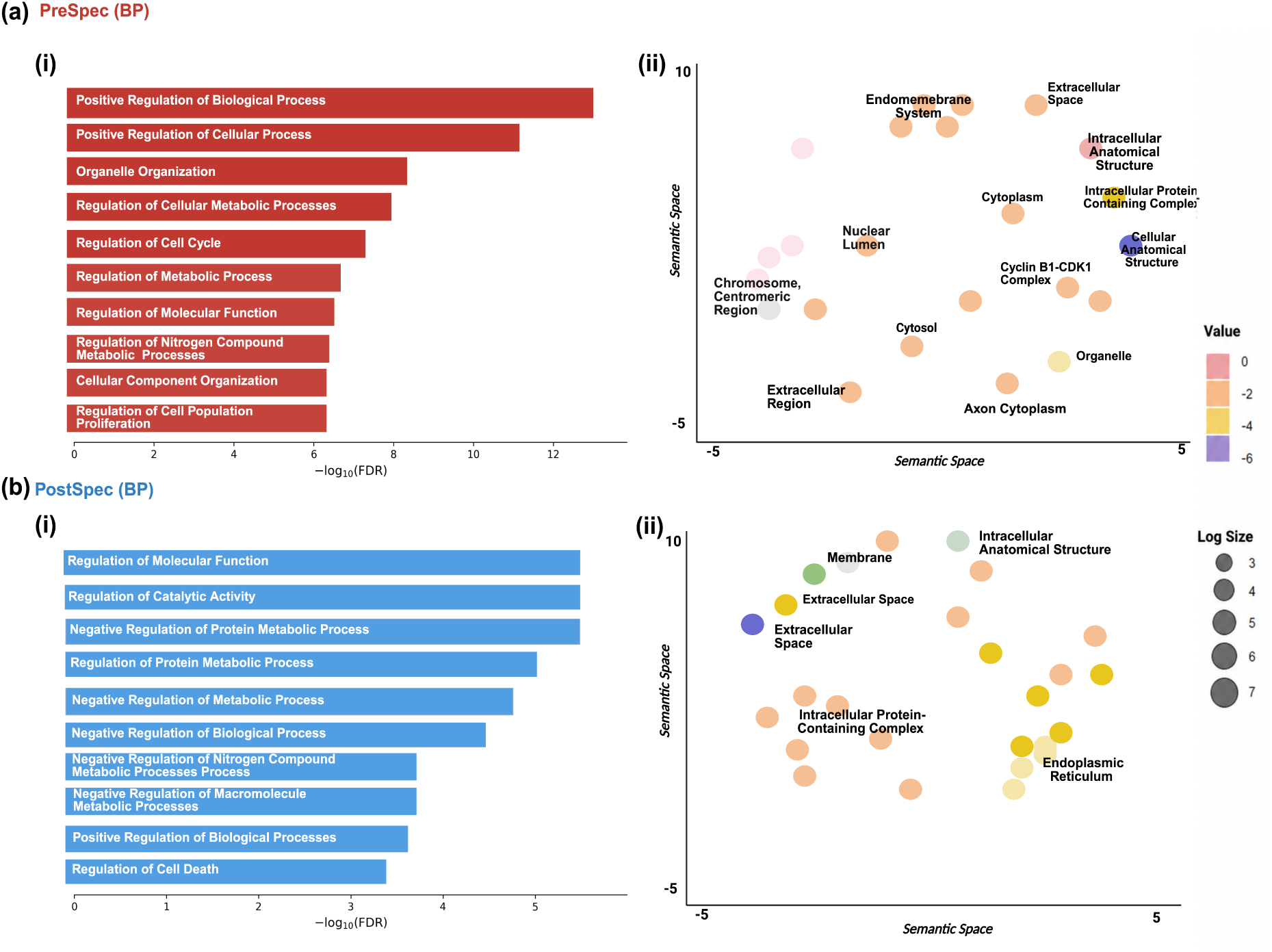
**(a)** (i) Top ten enriched Gene Ontology (GO) terms for PreSpec ohnologs Biological process terms are shown as a bar plot, and (ii) cellular component terms are shown as a bubble plot **(b)** (i) Top ten enriched GO terms for PostSpec ohnologs in paddlefish. (ii) Biological process terms are shown as a bar plot, and cellular component terms are shown as a bubble plot. Semantic similarity was calculated using REVIGO, and statistical significance was assessed using two-sided permutation tests (*p <* 0.01).

### Uncovering a missing duplicate *HoxC* cluster in the paddlefish assembly

The sterlet sturgeon genome has retained a particularly complete set of duplicate *Hox* clusters following the shared acipenseriform WGD. Paddlefish, while also having high retention of duplicate Hox clusters, have, according to two previous genome sequencing studies, only a single *HoxC* cluster, proposing that the other was secondarily lost after speciation (6, 12). These studies also inferred different total numbers of *Hox* genes in paddlefish, respectively 65 (6) and 75 (12), and as such we reassessed the Hox cluster and gene repertoire in our new duplicate resolved paddlefish assembly. We found a total of 77 *Hox* genes (Fig. 4 (c-d); Supplementary Materials (22)). Strikingly, we identified the missing *HoxC* cluster on chromosome 47 in our assembly which retained only one gene, *HoxC4*, suggesting that this cluster is highly degenerate. Notably, *HoxC4* is absent from the more intact *HoxC* cluster on chromosome 44 which is consistent with the previous chromosome-level assembly (12). Interestingly, *HoxC4* was identified in a previous scaffold level assembly (6), but placed within the non-degenerate cluster, meaning that as well as being at odds with our inferred *Hox* complement, the previous studies are at odds with each other (6). Furthermore, the *HoxC4* sequences identified here and in the scaffold-level assembly (6) are identical, indicating that there are likely assembly and/or annotation errors in at least one of the three genomes.

**Figure 4.**
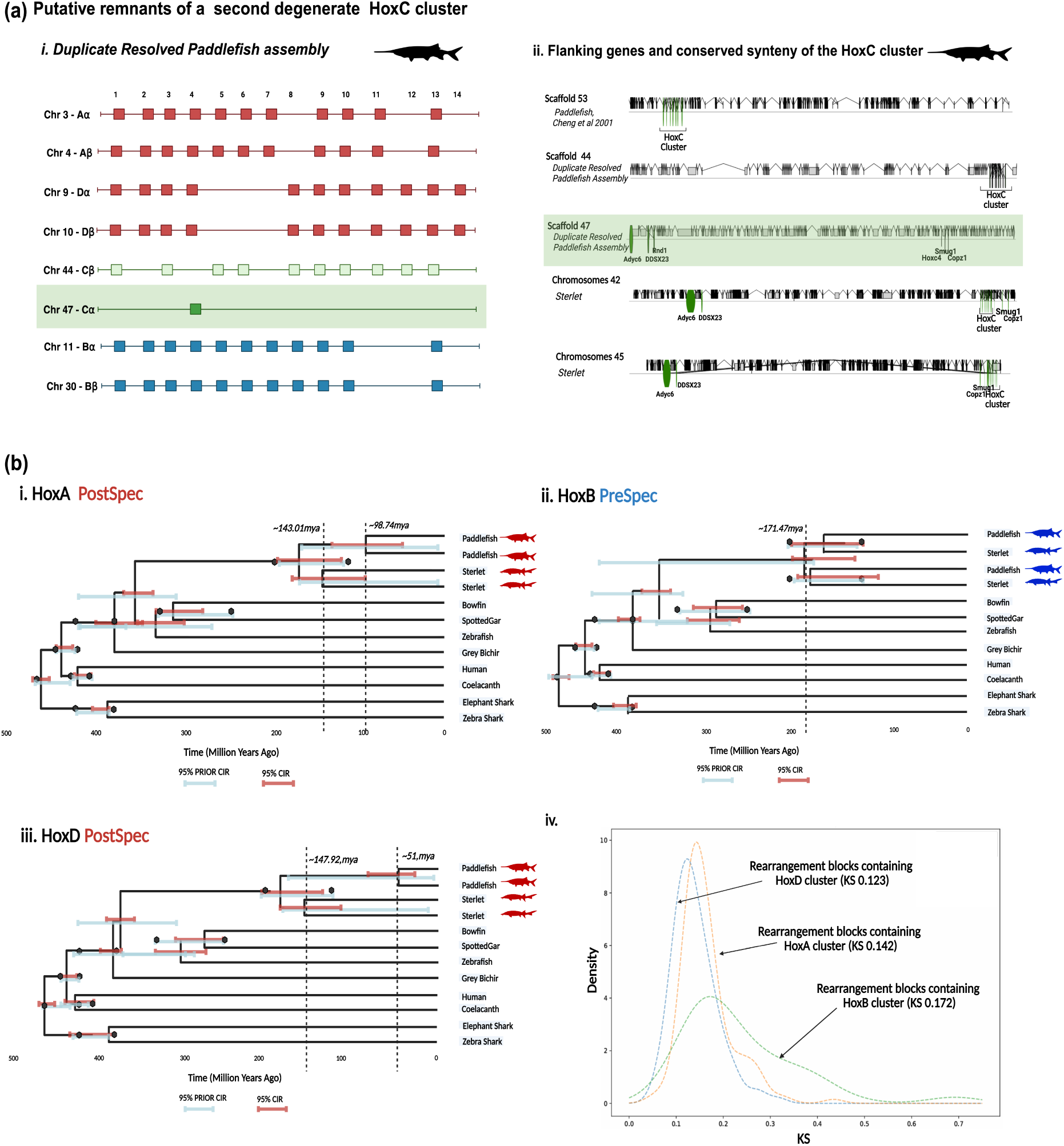
Hox cluster analysis in the Acipenseriformes. (a) (i) Hox genes in the duplicate-resolved paddlefish assembly, with remnants of a second *HoxC* cluster highlighted in green. (ii) *HoxC* cluster and flanking genes in the published paddlefish assembly, the improved assembly, and duplicated *HoxC* clusters in sterlet. The putative *HoxC* remnant in paddlefish, along with its flanking genes, is highlighted in green. (b) Dated phylogenies for (i) *HoxA*, (ii) *HoxB*, and (iii) *HoxD*. The 95% credibility interval (CIR) for each node is shown in red. Corresponding 95% CIRs from prior-only analyses are shown in blue below each node, demonstrating that WGD divergence time priors are sufficiently diffuse. Fossil calibrations (upper and lower bounds) are indicated by black diamonds. (iv) *K_s_* distributions for rearrangement blocks containing duplicated Hox clusters;*HoxA*, *HoxB*, and *HoxD* clusters in Paddlefish.

To rule out the possibility that the lone, degenerate-cluster-forming *HoxC4* gene on chromosome 47 of our new assembly was an artefact of misassembly or otherwise we first searched for genes commonly found to flank *HoxC* clusters; *tspan31, ddx23, rnd1, adcy6, smug1,* and *copz1* upstream and downstream of the intact *HoxC* cluster and the isolated *HoxC4* gene (91). None of these genes were found adjacent to the intact *HoxC* cluster but all expected flanking genes were found upstream or downstream of the single *HoxC4* on chromosome 47, and nowhere else in the genome (Fig 4 (a(ii)), Supplementary Materials (22)).

To exclude the possibility of misassembly, we also examined the read depth along chromosomes 47 and 44, identifying no abnormal signals in the regions containing the Hox clusters. Additionally, *HoxC4* was found in the scaffold-level assembly (6), located on scaffold JAAWVQ010136386.1, but all other *HoxC* genes were found as pairs on different scaffolds (*HoxC5* & *HoxC8* & *HoxC9* on scaffold JAAWVQ010005640.1 *HoxC11* & *HoxC10* on scaffold JAAWVQ010044606.1, *HoxC12* & *HoxC13* on scaffold JAAWVQ010102576.1). Furthermore, flanking single genes *smug1 and copz1* were located on the same scaffold as *HoxC4* in the scaffold assembly. As such, the scaffold-level assembly validates *HoxC4* as a lone remaining member of the the *HoxC* on chromosome 47 and does not support the inclusion of *HoxC4* within the more complete *HoxC* cluster It appears likely that, lacking chromosomal context, the gene was mistakenly assumed to belong to the intact cluster. Importantly, sequence alignment shows that *HoxC4* retains a conserved homeodomain, the canonical two-exon structure, and an intact YPWM hexapeptide motif, suggesting that it remains functional despite its isolated genomic location (Supplementary Fig. S1 (B)).

Collectively, these findings provide strong evidence for the loss of *HoxC4* from the cluster on chromosome 44 and suggest that the copy located on chromosome 47 represents the sole remnant of the otherwise missing duplicate cluster. Notably, *HoxC4* was one of many genes recovered through the genome enhancement performed here (Supplementary Materials; table 3(22)), highlighted the value added by this approach. These findings suggest that, despite being singletons distributed across two distinct chromosomes in paddlefish, both the full complement of *HoxC* and flanking genes may still function as they do in genomes where they remain clustered, a phenomenon that, to our knowledge has not previously been described in vertebrate *Hox* cluster evolution.

### Rediploidisation of duplicate *Hox* clusters

*Hox* clusters have been key in validating several WGD events including in the acipenseriformes (15, 77), in teleosts, as well as in ancient WGD events at the base of vertebrates (16, 25, 37). We reconstructed gene trees for each *Hox* ohnolog pair gene family but found that clear signal for their rediploidisation histories was difficult to resolve (Supplementary Materials (22)). *Hox* genes are short, highly conserved sequences that can often yield weak phylogenetic signal (80). To combat this, the concatenation method has been useful in resolving phylogenetic problems where weak signal obscures underlying evolutionary relationships, including to resolve the rediploidisation of salmonid *Hox* clusters (71). As *Hox* clusters are not interrupted by rearrangement events, and based on our above findings regarding the link with rediploidisation, we infer that the clusters rediploidsed as complete units and can safely and validly be analysed collectively using concatenation. By concatenating individual *Hox* genes along a cluster from species that span the vertebrate phylogeny and inferring phylogenetic trees we recover PostSpec topologies for *Hox* cluster pairs A and D, indicating that they rediploidised independently in sturgeon and paddlefish, and a PreSpec topology for *HoxB* clusters, indicating that they underwent shared rediploidisation prior to the divergence of sturgeon and paddlefish (Supplementary Fig S1). Rediploidisation timing of the concatenated *HoxC* duplicate clusters was less straightforward, as only one clear, intact cluster is present in paddlefish weakening the robustness of the timing estimate for this cluster. Nonetheless, we assessed the topology and found that the sterlet duplicates grouped together, consistent with a shared, independent duplication (rediploidisation), with the single paddlefish cluster sister to the sterlet pair.

To augment our results, we used the Approximately Unbiased (AU) test to assess for a statistically significant in the likelihood of the PreSpec and PostSpec topologies for each Hox cluster (72) (Supplementary Fig S1 (A) and Supplementary Materials; Table 4 (22)) Since AU tests can be applied to fixed topologies, they were also performed on individual *Hox* gene alignments, testing whether each alternative topology could be rejected. This supported the concatenation analysis: for *HoxA*, both PreSpec topologies were rejected in favour of PostSpec in 7/11 gene trees (*p <* 0.05), while the remaining 4/11 failed to reject any topology (inconclusive). For *HoxD*, both PreSpec topologies were rejected in 8/10 gene trees. *HoxB* showed the opposite pattern: the PostSpec topology was rejected in favour of PreSpec in 4/11 gene trees (*p <* 0.05), with the remaining 5/11 inconclusive (Supplementary Materials (22)). These findings indicate that *Hox* clusters rediploidised at markedly different time points, with *HoxB* rediploidising in the common ancestor of sterlet and paddlefish, whereas *HoxA*, *HoxD* and *HoxC* rediploidised independently in sterlet and paddlefish.

### Phylogenomic dating of the *Hox* cluster rediploidisation

Redmond et al (2023), estimated the timing of the shared sturgeon–paddlefish WGD using ohnologs that rediploidised prior to species divergence, providing mean lower-bound estimates for the WGD event at approximately *∼* 254.7 Ma and *∼* 241.8 Ma (with joint 95% confidence intervals spanning 202.9-289 Ma), depending on which fossil calibrations were applied. They also estimated the speciation event splitting sturgeons and paddlefish at *∼* 171.6 Ma or *∼* 167.5 Ma depending on calibration strategy, with joint 95% confidence intervals spanning 123.4-203.3 Ma, suggesting a Jurassic or perhaps early Cretaceous divergence consistent with many other studies (70). This estimate is substantially different to previous estimates which lacked the asynchronous rediploidisation context (12, 15) Here we estimate divergence times of the concatenated *Hox* gene datasets, expecting a long-lag time post-WGD (given the time between the WGD and sturgeon-paddlefish lineage divergence) and anticipating the possibility of vastly different times in each species for the lineage-specific rediploidisation of the *HoxA* and *HoxD* clusters.

For *HoxA*, which rediploidised after speciation, we infer the timing of independent rediploidisation events: in paddlefish at *∼* 98.74 ma (95% CI: 137.47–60.01 Ma), and in sturgeon at *∼* 143.01 ma (95% CI: 182.93–103.10 Ma), suggesting these events may have occured at very different times, although the confidence intervals overlap. *HoxD* also follows a post-speciation rediploidisation scenario, but with a clear and striking temporal difference: in paddlefish at *∼* 51 Ma (95% CI: 80.27–23.14 Ma), and in sturgeon at *∼* 147.92 ma (95% CI: 182.84–113.00 Ma). *HoxB*, on the other hand, rediploidised in the ancestor of sturgeon and paddlefish, with an inferred rediploidisation time of *∼* 171.57 ma (95% CI: 145.59–212.51 Ma), perhaps coming only very shortly before their lineage divergence (Fig 4). These findings present a clear picture of asynchronous rediploidisation occurring at *Hox* clusters, with stark temporal differences that may reflect the distinct biological and morphological differences observed between the two lineages. To connect the rearrangement blocks defined in the previous section to the *Hox* cluster dating, we assessed whether these blocks containing the clusters could themselves be “dated” using *K_s_*, and whether the signal was consistent with the more precisely resolved timing of the enclosed *Hox* cluster. In Fig. 4 (b)(iv) and Supplementary Fig. S1 (c), the blocks containing *HoxA* and *HoxD* show *K_s_* peaks close to zero, consistent with more recent rediploidisation and correspondingly fewer subsequent substitutions. The block containing *HoxB*, by contrast, shows a broader peak shifted further from zero, consistent with an older rediploidisation event. This is broadly consistent with the *K_s_* analysis reported in previous studies and supports our conclusion that blocks of genes rediploidise together via structural rearrangements (70).

## Discussion

Rearrangement events are often presumed as the mechanism by which rediploidisation occurs following an autopolyploid WGD event, yet only a handful of studies have detected patterns that support this. Here, we have directly taken advantage of the signal coming from asynchronous rediploidisation in slowly-evolving sturgeon and paddlefish genomes i.e. syntenic clustering of genes with shared rediploidisation history (duplication times), to test this. We identify significant associations between genome rearrangement blocks and gene tree topology duplication time blocks, providing strong evidence that rearrangements are key mechanistic drivers of rediploidisation.

We present an updated and enhanced genome assembly for the American paddlefish adding 9,859 genes to the genome. Remarkably, despite the WGD event being estimated to have occurred a quarter of a billion years ago, large portions of this genome remain highly similar and were consequently prone to assembly collapse. Here, we present a method for improving the assembly, together with our enhanced annotation. While sequencing techniques have advanced and long-read sequencing is much more commonplace, we highlight the implications for future studies, cautioning that similar assembly errors are likely to affect not only recent polyploids but also more ancient events as exemplifed here (79). Epitomizing the value of our improved assembly and annotation we recovered two HoxC cluster regions in paddlefish for the first time after multiple previous studies indicated that one copy had been lost. The newly identifed cluster is degenerate and consists of only a HoxC4 gene surrounded by known HoxC flanking genes. Interestingly our analysis of the Hox cluster rediploidsation patterns supports independent, lineage-specific rediploidsation of the HoxC clusters in sturgeon and paddlefish. This implies that the Hox cluster duplicated independently in each lineage, and consequently the functional trajectories of each cluster pair are expected to be entirely lineage specific. It is therefore important to emphasize that the genetic and functional state of the sturgeon cluster cannot be interpreted as the ancestral condition of the more degenerate paddlefish cluster pair.

Understanding how different ploidy states can persist in a stratified manner along chromosomes over tens of millions of years requires a deeper understanding of meiotic processes following WGD. One potential additional factor is higher-order genome organisation The association we observe between chromosomal rearrangements and TAD boundaries suggests that the 3D genome architecture may influence where and how rediploidisation occurs, with genes located within the same TAD tending to rediploidise together. If TAD boundaries and rediploidisation timing are indeed linked, this would imply that disruption of TAD structure is selectively disadvantageous. In this framework, TADs may act as both structural and functional constraints on the rediploidisation process by restricting the range of viable chromosomal rearrangements. Rearrangements that disrupt established regulatory domains could be deleterious, whereas those that preserve or accommodate existing TAD architecture would be more likely to avoid purifying selection Under such a model, the maintenance of regulatory integrity and its complexity might not only be expected to constrain the space and patterns of rediploidisation possible but to explain, at least in part, the asynchronous nature of rediploidisation observed here and in other lineages (50, 65, 71, 90). Alternative explanations are also possible however Rather than constraining rediploidisation, TAD architecture may itself be shaped by the rediploidisation process, or it may even be a complex blend of both, across genomic regions Distinguishing between these possibilities will require further investigation. Nevertheless our findings suggest that 3D genome organisation may represent an important, and largely unexplored, component of the evolutionary dynamics of post-WGD genomes.

Our analyses show that early-rediploidising regions are enriched for pathways involved meiotic regulation, cell-cycle control, chromosome segregation, DNA repair, genomic stability, and stress resistance. By contrast, lineage-specific rediploidising regions show little evidence of functional enrichment, suggesting that they are functionally more heterogeneous. This finding ameliorates findings in salmonids and *Arabidopsis arenosa* in which ohnologs associated with cell-cycle regulation and genome maintenance exhibit signatures of adaptive evolution and selection following WGD (27, 34, 55). Although these studies demonstrate that such WGD-derived genes can be targets of adaptive evolution, our results suggest a more specific pattern: genes associated with cell-cycle regulation and genome maintenance are disproportionately represented among early-rediploidising regions This enrichment is consistent with a model in which the earliest phase of rediploidisation contributed to the stabilisation of cellular functions required for chromosome pairing, segregation fidelity, and genome integrity following WGD. More broadly, our findings raise the possibility that particular functional classes of genes are preferentially associated with early rediploidisation.

*Hox* clusters provide clear examples of genomic regions that rediploidised at different times with cases found both before and after the paddlefish-sturgeon divergence. Our phylogenetic analyses, together with divergence time estimates, strongly support a highly asynchronous pattern of rediploidisation among the clusters. As discussed in (70), divergence estimates that fail to account for asynchronous rediploidisation are likely biased toward more recent dates because they instead capture the timing of average, or most recent ohnolog divergence rather than the WGD event itself. Previous attempts to date the paddlefish WGD event were probably affected by this, such as an inferred date for an independent WGD event in the paddlefish lineage at approximately *∼* 42 Ma based on sequences from the *HoxA* and *HoxD* clusters (15). Our results show that this estimate instead closely corresponds to the timing of rediploidisation within the paddlefish *HoxD* cluster. Correctly accounting for these processes is therefore essential for reconstructing the history and timing of WGD (13, 53, 68, 70).

*Hox* genes are central regulators of body patterning, making their independent rediploidisation particularly noteworthy in the context of lineage-specific morphological evolution (32). We estimate that the *HoxD* cluster in paddlefish rediploidised relatively recently, at approximately *∼* 50 Ma as a mean estimate, whereas the corresponding region in sterlet rediploidised much earlier, around *∼* 147 Ma. This near 100-year mean difference in duplication time of such an iconic gene cluster provides a stark demonstration of the lineage-specific trajectories rediploidisation can take if it has not completed before speciation (28). Furthermore,assuming rediploidisation was underway at *∼* 254 Ma (70), this implies that some regions, including this HoxD cluster pair, of the paddlefish genome may have remained effectively tetraploid, with duplicate genes not resolving from tetrasomic singletons, for over 200 million years (as a mean estimate). *HoxA*, *HoxD* and *HoxC* all appear to show rediploididsation post-speciation and thus provide a striking example of extreme asynchronous rediploidisation, with homologous genomic regions resolving to diploid inheritance only after remarkable delay and on vastly different evolutionary timescales in the two lineages. Such differences in gene duplication status may have contributed to the distinct morphologies observed in paddlefish and sturgeons given the role of *Hox* genes in development (32, 33). For example, paddlefish is characterised by a dramatically elongated rostrum rich in electroreceptors, an adaptation for detecting planktonic prey, while sturgeons have a much shorter snout with sensory barbels and are adapted for benthic suction feeding. Whether delayed rediploidisation of *Hox* clusters contributed to these and other morphological differences between the lineages is of course unclear, but warrants further functional exploration. Interestingly, the estimated timing of rediploidisation for the *HoxB* cluster falls close to the inferred paddlefish–sturgeon divergence time (70), which places the split at approximately *∼* 171.6 Ma or *∼* 167.5 Ma This suggests that the *HoxB* cluster likely rediploidised shortly before speciation. Different rediploidisation timings of *Hox* clusters is not unique to Acipenseriformes and has also been documented for *Hox* duplicates in salmonids and other teleosts (56, 71).

Our findings evidence genomic rearrangements as the mechanism of rediploidisation, and further support it as an asynchronous and prolonged evolutionary process that generates ohnologs, sometimes long after WGD. Conflating rediploidisation with the WGD event itself, or using rediploidisation (gene divergence and its timing) as a proxy for WGD, is potentially highly problematic, particularly in autopolyploids (5, 50, 65, 70, 71) Reassessing ancient WGDs with consideration for the mode and tempo of rediploidisation may therefore reshape our interpreation of polyploidisations, including for key events such as those in early vertebrate evolution.

## Materials and Methods

### Resolving collapsed duplicates in the paddlefish genome

Cheng et al. (2021)’s American Paddlefish assembly has 60 pairs of chromosomes with a genome size of 1.54GB with 26,017 predicted protein-encoding genes (12). The assembly was sequenced to 30X coverage and short and long reads from this study were downloaded from CNGB under project accession number CNP0000867. Cheng et al. (12) note a smaller than expected genome size following a 17-mer analysis (51). Following a similar method to Du et al.(17), we identified these collapsed regions and attempted to resolve them (39, 42, 93).

PacBio long-reads were aligned with bwa (v0.7.17-r1198-dirty) (48) with default parameters. This was sorted and indexed using SAMtools (v1.16.1) (8). The short reads were aligned with Bowtie (v2.4.2) (45) using standard parameters and again, indexing and sorting was done with SAMtools (v1.16.1) (8). To assess the depth of coverage across the genome, the alignments were split into 10kb regions and mosdepth (v0.3.3) (66) was used to quantify read depth at each segment with parameter *–by*. 10kb regions with double the expected read depth (double coverage segments), were separated from the rest of the genome for the next steps. Using FreeBayes (v1.3.6) (26), a polymorphism VCF was generated from the short-read alignments. After this, long-reads were used to separate “haplotypes” in the double coverage regions. HapCUT2 (20), a haplotype assembly tool, was used to reconstruct individual haplotypes from double-coverage long-read alignments. This approach enables the identification of genomic regions containing more than one haplotype and was thus used here to detect potential duplicated loci represented mistakenly as multiple alternative alleles in the double-coverage long-read BAM file. HapCUT2 requires as input a BAM file of mapped reads and a corresponding VCF file (20). The software is designed for diploid genomes and does not currently support phasing of polyploids Parts of the paddlefish genome however, exhibited signatures of apparent polyploidy To overcome the limitations of HapCUT2 we filtered the VCF to remove polyploid or erroneous genotype calls (e.g., 4/4, 3/4) prior to analysis. Polyploid genotype fields (e.g. 4/4) were coerced to diploid genotypes (2/2) using a custom script. While this does not fully capture the underlying ploidy, it preserves more information than discarding such variants entirely by retaining the two most common alternative alleles. HapCUT2 results for our double-coverage regions contained multiple assembled haplotypic segments for a given loci (collapsed duplicates) which were subsequently split into individual files and annotated with corresponding VCF information using a custom script by Du et al (17) (available at https://github.com/dukecomeback/sterletM sch) that was modified for this study. These split files were then processed with fgbio’s *HapCutToVcf* script to generate separate VCF’s for each assembled haplotype. These VCF files and the reference were used to produce haplotypic contigs in fasta format using *vcf-consensus* from the bcftools package (v1.10.2) (47). The fasta files of the split regions were merged with the original contigs (minus the double coverage contigs) using the Unix *cat* command (See Supplementary Figure 1).

### Assembly and scaffolding of the paddlefish genome

Following haplotype splitting, the genome was then reassembled and scaffolded with the available HiC data. Cheng et al. (12) described 60 pairs of chromosomes (n=120), finding 26 macro chromosomes and 34 smaller, micro-chromosomes, a number which aligns with previous karyotype studies (77) and is equivalent to the sterlet genome (17). Assembly and scaffolding were done using Juicer (v1.6) and 3D-DNA (v190716)(18, 19). The Illumina short-reads were aligned to the duplicate resolved contigs with Juicer (v1.6)(19). 3D-DNA (v190716) (18) was then used for assembling the genome with -r=0 flag to ensure no iterative rounds of mis-join correction were carried out. Finally, the scaffold assembly was manually reviewed using Juicebox assembly tools (v1.6)(18).

### Paddlefish RNA-seq and genome annotation

To produce high-quality gene predictions we first produced RNA-seq data for fifteen tissues from four paddlefish individuals. Captive-bred American paddlefish were purchased from Osage Catfisheries Inc. (Osage Beach, MO, USA) and euthanized on site by MS-222 overdose, with samples collected from brain, caecum, eye, gill filament, gill raker, gonad heart, kidney, liver, rostrum, skeletal muscle, skin, spiral valve, spleen, and stomach from four individual fish. Samples were flash frozen using liquid nitrogen and stored in RNAlater at -80°C. Total RNA was prepared by lysing tissue samples in 1ml Tri Reagent (Sigma Aldrich) using a Tissue lyser LT (Qiagen) and 7mm steel bead. Following incubation for 5 minutes at room temperature, 0.2ml of chloroform was added and the tube shaken for 15 seconds. After incubation for 2 to 3 minutes at room temperature the tubes were spun at 12,000g for 15 minutes. The upper, RNA-containing aqueous fraction was moved to a clean 1.5ml tube, mixed with an equal volume of 70% ethanol, and vortexed This solution was added to a spin column from the RNeasy mini kit (Qiagen) and RNA prepared following the manufacturer’s instructions, including the DNaseI treatment step to remove any contaminating genomic DNA. Purified RNA was quantified using a Qubit 3.0 Fluorometer and then stored at -80°C. Quality control, poly-A enrichment mRNA library preparation and RNA sequencing of 30 million 150bp paired end Illumina reads were performed by Novogene America (Sacramento, CA, USA). Summary of paired-end RNA-seq FASTQ files generated from *Polyodon spathula* (paddlefish) tissues are shown in Table S4 and have been deposited in the NCBI Sequence Read Archive (SRA) under BioProject accession PRJNA1512476.

Genome annotation was performed using BRAKER3 (30), integrating de novo gene prediction with protein homology and RNA-seq evidence. Assembly completeness was assessed using BUSCO (v5.4.4) (54) with the Actinopterygii odb9 dataset, run with the-augustus and -long options enabled (54, 75). Protein homology evidence was provided to BRAKER3 using the Metazoa protein database together with proteomes from the following selected vertebrate genomes, used also in orthology inference (12): American paddlefish, elephant shark, zebrafish, medaka, fugu, stickleback, sea lamprey, spotted gar human, mouse, and sterlet. Protein sequences were clustered using CD-HIT (24) and supplied to BRAKER3 using the --prot seq option. Raw read quality was assessed using FastQC (v0.11.9) (2) . Reads were trimmed to remove adapters and low-quality bases using Trimmomatic (v0.39) (7). RNA-seq reads from multiple adult paddlefish tissues were aligned to the genome using HISAT2 (v2.2.1) (40). RNA-seq reads from multiple adult paddlefish tissues were aligned to the genome using HISAT2 (v2.2.1) (40). Alignments were sorted and indexed with SAMtools (v1.16.1) (8) and provided to BRAKER3 via the --bam option. BRAKER3 was run in combined RNA-seq and protein mode using GeneMark-ETP and AUGUSTUS for gene prediction, enabling automated training and evidence integration without manual model reconciliation. Functional annotation of predicted gene models was performed using the FANTASIA pipeline (57).

### Orthology Assignment

Orthofinder (v2.5.4) (21) was used for orthology inference. We included a diverse set of proteomes that spanned the jawed vertebrate phylogeny including from the Chondrichthyes: ghost shark (*Callorhinchus milii* ; GCF 000165045.1) (85); from the Sarcopterygii; human (*Homo sapiens*; GCF 000001405.39), African clawed frog (*Xenopus tropicalis*; GCF 000004195.4), and coelacanth (*Latimeria chalumnae*; GCF 000225785.1) from within Actinopterygii we selected zebrafish (*Danio rerio*; GCF 000002035.6), spotted gar (*Lepisosteus oculeatus*; GCF 000242695.1) (9), Grey bichir (*Polypterus senegalus*; GCF 016835505.1) (6). We included a species tree in our Orthofinder (21) run with flag *-s* in line with the accepted relationships in the jawed vertebrate phylogeny to augment orthology inference:*((Callorhinchus milii), ((Latimeria chalumnae,(Xenopus tropicalis,(Homo sapiens)),(Grey birchir,((Polyodon Spathula, Acipenser ruthenus),((Danio rerio), (Lepisosteus oculatus)))))))*.

We extracted the longest isoform from each proteome using a custom pipeline, reduced sequence redundancy with CD-HIT (24), and then ran OrthoFinder (v2.5.4) (21) using the -s parameter to include a rooted species tree. To improve the resolution of orthogroup relationships, we additionally used the -y flag in OrthoFinder when generating Phylogenetic Hierarchical Orthogroups (PHOGs). This option separates paralogous clades below the root of a HOG into distinct hierarchical orthogroups, thereby reducing the lumping of multiple sturgeon–paddlefish ohnolog pairs into single gene families during downstream analyses. Following the protocol described in Redmond et al. (70), PHOGs were further filtered by extracting groups containing two sequences each from sturgeon and paddlefish, together with at least one outgroup sequence for subsequent rooting of the sturgeon–paddlefish subtree. Sturgeon and paddlefish genes were only selected from the 60 largest chromosomes.

### Ohnolog duplication time inference

The sturgeon-paddlefish ohnolog pairs described above, were subjected to phylogenetic analysis to estimate the time of rediploidisation relative to the speciation event. MAFFT (v7.453) was used for multiple sequence alignment of the filtered PHOGs with standard parameters. Phylogenetic inference by ML was performed with IQ-TREE (v1.6.12)(60) with the -m JTT+G flag, -bb 1000 flag allowing 1000 ultrafast bootstrap (59) replicates These ohnolog gene trees were processed and analysed for duplication time inference (i.e rediploidisation time). Scripts described and published in (70) were used to infer the rediploidisation time of each gene tree. The resulting gene trees were summarised into different groups indicating their duplication time: PreSpec, PostSpec, Other[PreSpec-like PostSpec-like].

### Hox cluster analysis

We downloaded the complete Hox cluster sequences from the spotted gar (9), sterlet (17) zebrashark (91) and elephant shark (85). TBLASTN (NCBI BLAST+ suite v2.17.0) was used to align these genes against our genome assembly. Taking the longest captured regions we then used Exonerate (v2.4.0)) to align the best-hits (73) and delineate and verify the open-reading frames of 77 Hox genes across 8 chromosomes. We also took the protein sequences of elephant shark and human *HoxC* flanking genes *Tspan31*, *Ddx23*, *Rnd1*, *Adcy6*, *Smug1*, and *Copz1* to verify the location of duplicate *HoxC* clusters. To investigate the phylogenetic relationships of the clusters, we used *Hox* orthologs from *Acipenser ruthenus* (sterlet), *Polypterus senegalus* (gray bichir), *Amia calva* (bowfin) *Stegostoma tigrinum (zebrashark)*, *Callorhinchus milii* (elephant shark), *Danio rerio* (zebrafish), *Lepisosteus oculatus* (spotted gar), *Homo sapiens* (human), and *Latimeria chalumnae* (coelacanth), obtained from the NCBI database. Sequences were aligned using MASCEv2 (69) and codon alignments were generated using the amino acid alignments as a templates with the same software. These alignments were used to construct ML gene trees for each *Hox* gene in IQTREE (v.2.1.3). using ModelFinder, for model selection and the -bb 1000 flag allowing 1000 ultrafast bootstrap replicates(UFBOOT). For the concatenated alignments the best-fitting model for each concatenated gene family was again determined using IQ-TREE (v.2.1.3). To provide statistical delineate between these classifications, the Approximately Unbiased (AU) test was employed in IQ-TREE (v.2.1.3) to compare the likelihood of the two possible PresSpec against the single PostSpec topology. Gene families were assigned to the best-supported topology where alternative hypotheses could be rejected at *p <* 0.05.

### Defining Micro-and Macro-Synteny blocks and Topology Break**points**

Micro-syntenic blocks between the sturgeon and paddlefish genomes were identified using OrthoFinder (v2.5.4) (21) and i-ADHoRe (v3.0.01) (67). Proteomes from sterlet, the duplicate-resolved paddlefish assembly, and gray bichir were analysed using OrthoFinder with default parameters. Conserved syntenic blocks were subsequently identified with i ADHoRe, and genomic coordinates for genes within each block, as well as block boundaries (defined by the first and last gene pairs), were extracted from the segments.txt output file. Macro-synteny blocks, representing larger-scale rearrangements, were constructed following the methodology described in (63) To identify genomic regions associated with transitions in ohnolog gene-tree topology, a Bayesian segmentation approach was applied to paddlefish and sturgeon chromosomes using the algorithm available at https://github.com/McLysaght-Evolutionary-Genetics/bayesian-segmentation. In this approach, each chromosome was modeled as an ordered sequence of categorical observations corresponding to posterior probabilities of the gene-tree topology states: PreSpec, PostSpec A topology breakpoint was defined as the genomic position at which the most probable topology state transitioned between categories. To assess spatial relationships between chromosomal rearrangements and changes in rediploisation history, genomic distances were calculated between each micro-and macro-synteny block boundary and the nearest topology breakpoints. These observed distances were compared against null distributions generated from randomly sampled genomic coordinates matched by chromosome and block size. All distance calculations and randomisations were performed using custom Python scripts (86);(Supplementary Materials (22)). PreSpec and PostSpec gene trees within each synteny block were quantified using custom scripts (Supplementary Materials (22)). To test whether specific gene categories (PreSpec or PostSpec) were overrepresented within individual synteny blocks relative to genome-wide background frequencies, Fisher’s exact tests were performed. For each block and category, a 2×2 contingency table was constructed comparing the number of genes of the focal category within the block to the number outside the block. Statistical analyses were conducted using the *scipy* Python package, with significance evaluated at *p <* 0.05.

### Functional investigation with STRING

Protein–protein interaction networks were analysed using STRING v12.0. (78) All predicted protein sequences were queried against the STRING database with the American Paddlefish proteome used as background. STRING’s PPI enrichment test was used to evaluate whether observed interactions exceeded random expectation given the size of each protein set. A PPI enrichment p-value *p <* 0.05 was considered indicative of statistically significant interaction structure. Functional enrichment assessments were conducted using STRING’s integrated annotation tools. For each gene set, enrichment analyses were performed across:Gene Ontology (Biological Process, Molecular Function, Cellular Component), KEGG pathways, Reactome pathways and STRING Local Network Clusters (LNCs). Significance was evaluated using the Benjamini–Hochberg false-discovery rate (FDR) method, with *FDR <* 0.05 considered significant. STRING’s default species-wide proteome was used as the background set for all tests. Semantic similarity was calculated using REVIGO, and statistical significance was assessed using two-sided permutation tests (*p <* 0.01).

### TAD boundaries

Hi-C data were processed using the HiCExplorer v3.7.6 pipeline (89). Raw Hi-C reads were aligned to the reference genome using bamTools (3), and the resulting BAM files were sorted and filtered before matrix construction with hicBuildMatrix with the follow ing parameters: –binSize 10000 –restrictionSequence GATC (mbol) –danglingSequence GATC –minDistance 150. The Hi-C matrix was normalised using Knight–Ruiz (KR) balancing with hicCorrectMatrix to correct for coverage biases. TADs were identified using hicFindTADs with FDR correction and the following parameters: *–minDepth 3000 –maxDepth 31500 –minBoundaryDistance 30000 –step 1500 –thresholdComparisons 0.005 –delta 0.01 –correctForMultipleTesting fdr.* These analyses were also attempted for the sterlet sturgeon, but at least with currently available data we were unable to robustly identify TADs and their boundaries (data not shown).

### Phylogeneomic divergence dating of hox clusters

Phylogenomic divergence dating was conducted using PhyloBayes v4.1 (46), applying the site-heterogeneous CAT-GTR+G4 substitution model with an autocorrelated log-normal relaxed clock (36), and a birth-death prior with soft fossil calibration bounds (35, 92). Fossil calibration priors (Supplementary Materials (22)) were applied to most nodes in the tree, with the exception of the sturgeon–paddlefish WGD lower-bound node and the node separating Acipenseriformes (sturgeons and paddlefish) from Neopterygii The calibration scheme largely followed that of (70), which is based on (4). Specifically, a minimum divergence of 121 Ma for sturgeons and paddlefish was used (76), and a lower bound of 381 Ma was set for crown Chondrichthyes (23). A fixed tree topology was specified based on established jawed vertebrate phylogeny(12, 17, 36, 70) and our inference of a shared WGD. The topology used for *Hox* A and *Hox* D was: ((zebra shark,elephant shark),((coelcanth,human),(grey bichir,((zebrafish,(spotted gar bowfin)),((sterlet,sterlet1),(paddlefish,paddlefish1)))))); was specified based on accepted jawed vertebrate phylogeny (12, 17, 36) Zebra shark and whale shark (Chondrichthyes) were designated as outgroups. To confirm this topology, we conducted a basic concatenated phylogenomic analysis for each of our five datasets using IQ-TREE (60) under the LG+G4 model, with 1000 UFBoot bootstrap replicates. Each PhyloBayes MCMC chain was run for at least 30,000 cycles, discarding the first 5,000 as burn-in before calculating divergence times and 95% credibility intervals (CIRs). To verify the influence of priors, we also performed runs under the prior using the same settings but substituted the site-heterogeneous CAT-GTR+G4 model with a site-homogeneous Poisson model for computational efficiency. Because the prior over divergence times is independent of the substitution model, this approach allowed us to confirm that our WGD divergence priors were sufficiently diffuse and did not unduly constrain the main analysis.

### *K_S_* analysis of the *Hox* clusters

Synonymous substitution rates (*K_S_*) values were calculated for *Hox* gene clusters in paddlefish and sterlet. In addition, we included paralogous gene pairs from within rearrangement blocks containing the *Hox* clusters in each species. We assume that these blocks consist of genes that experienced rediploidisation contemporaneously with the Hox clusters. *K_S_* values were estimated for each rearrangement block containing a Hox cluster pair using the wgd v2 toolkit (11) for each species.

## Acknowledgments

This work was supported by funding from the European Research Council, grant agreement 771419 (to A.McL.). A.K.R. was supported by an Irish Research Council Government of Ireland Postdoctoral Fellowship (GOIPD/2021/466), a Royal Society-Research Ireland University Research Fellowship (URF*\*R1*\*241884), and a UCD Ad Astra Fellowship D.J.M. was supported by the Biotechnology and Biological Sciences Research Council through grants BBS/E/RL/230001B and BB/Z51746X/1

## Supplementary Figures and Tables

**Table S1.** Categorisation of the possible rooted sturgeon-paddlefish subtrees with duplication nodes coming before (‘PreSpec’) or after (‘PostSpec’) the species diverged and ‘Other’ trees that only partially match one of these scenarios (either, ‘PreSpec-like’, or ‘PostSpec-like’). Counts from Redmond et al 2023 and this study shown here

|  | PreSpec<br>Trees | PostSpec<br>Trees | Other<br>Trees |
| --- | --- | --- | --- |
| Redmond et al. 2023 | 1448 | 2074 | 1917 |
| This study | 1713 | 2105 | 2044 |

**Table S2.**
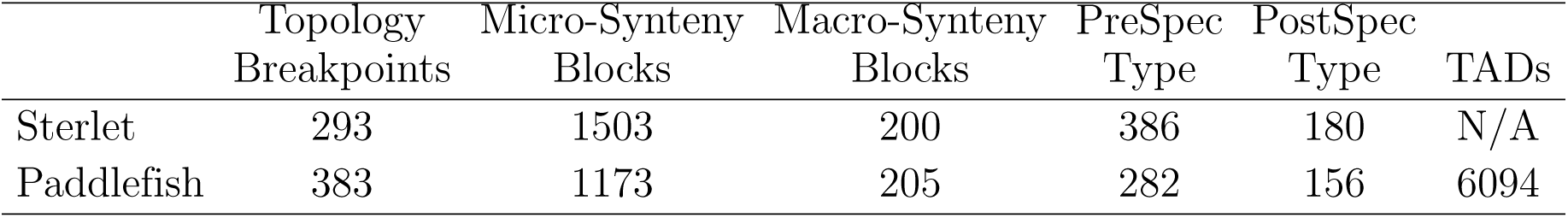
Counts of the different breakpoint types identified in the study, as well as counts for PreSpec-type and PostSpec-type blocks. Blocks were classified as PreSpec-type or PostSpec-type when they showed a significant enrichment of PreSpec or PostSpec genes relative to the genomic background (one-sided Fisher’s exact test, *p <* 0.05).

**Table S3.** Assembly stats.

|  | Paddlefish, Cheng et al. 2021 | Paddlefish, Duplicate Resolved Assembly |
| --- | --- | --- |
| Genome Size | 1.54GB | 1.56GB |
| Scaffold N50 | 48.9MB | 48.9GB |
| GC content | 39% | 39.2% |
| Protein-Coding Genes | 26,071 | 35,930 |
| Chromosome No. | 60 pairs (2n+120) | 60 pairs (2n+120) |
| BUSCO | 93%[S:51%,D:41%],F:2.5%,M:5.5%,n:3640] | 93.14[S:36.5%,D:56.6%],F:1.3%,M:5.6%,n:3640] |

**Table S4.**
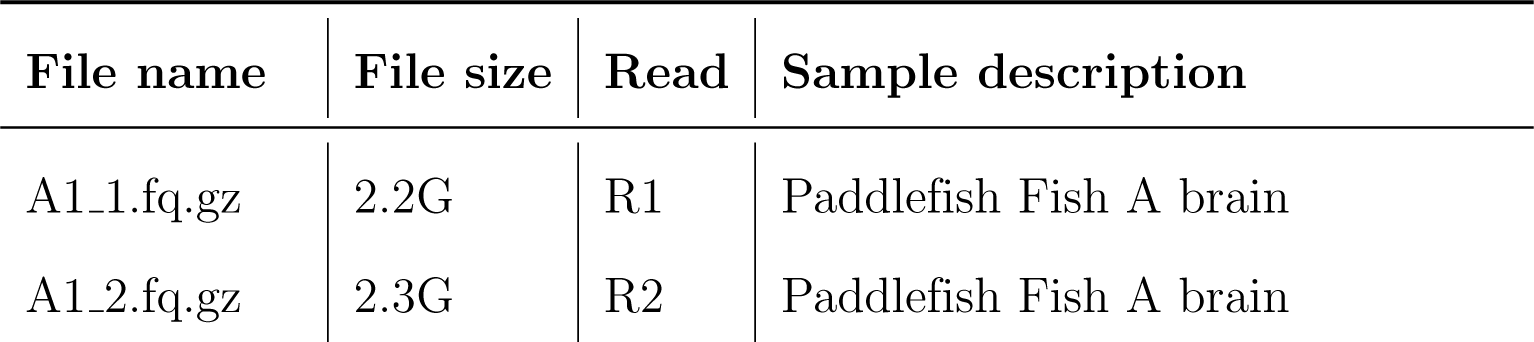
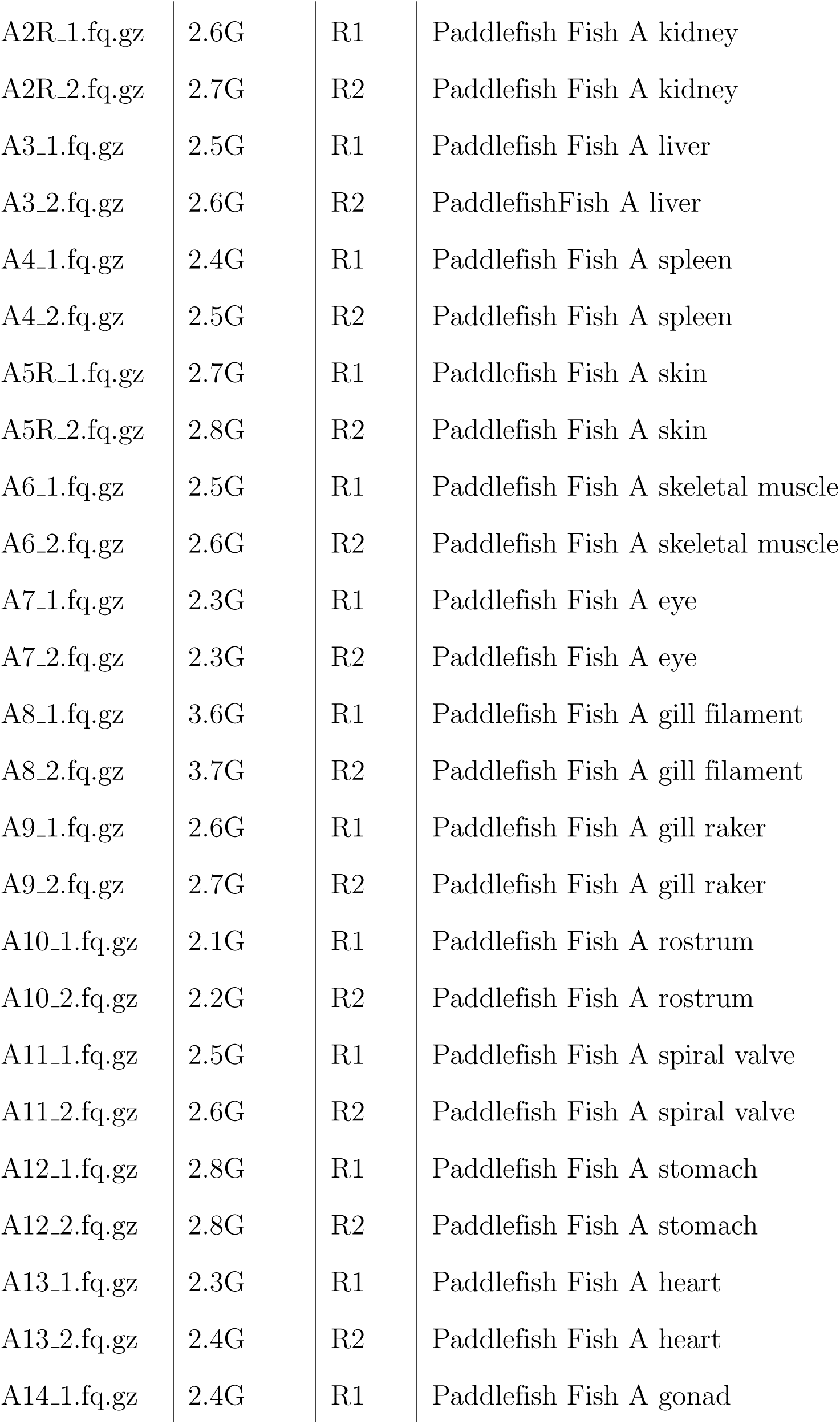
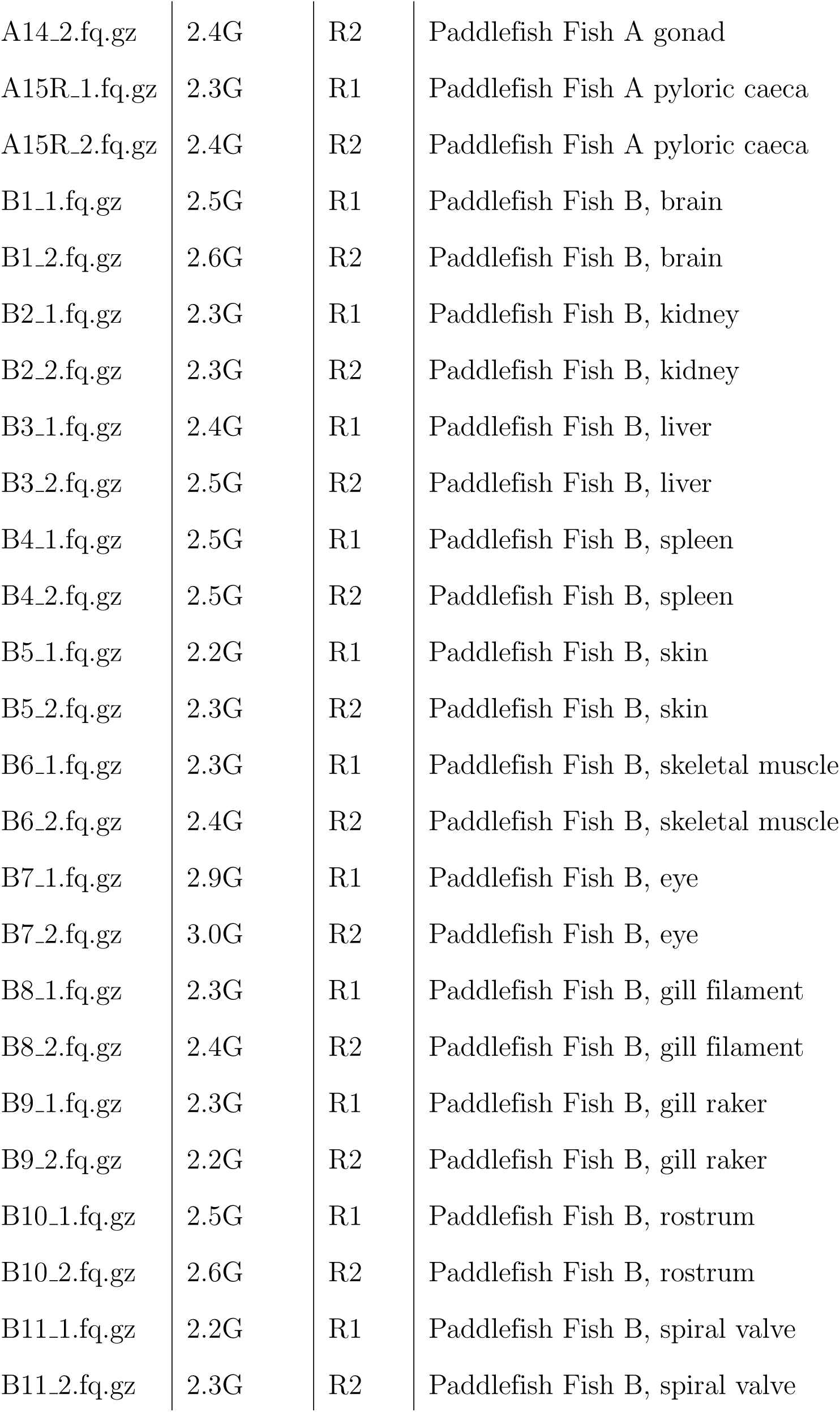
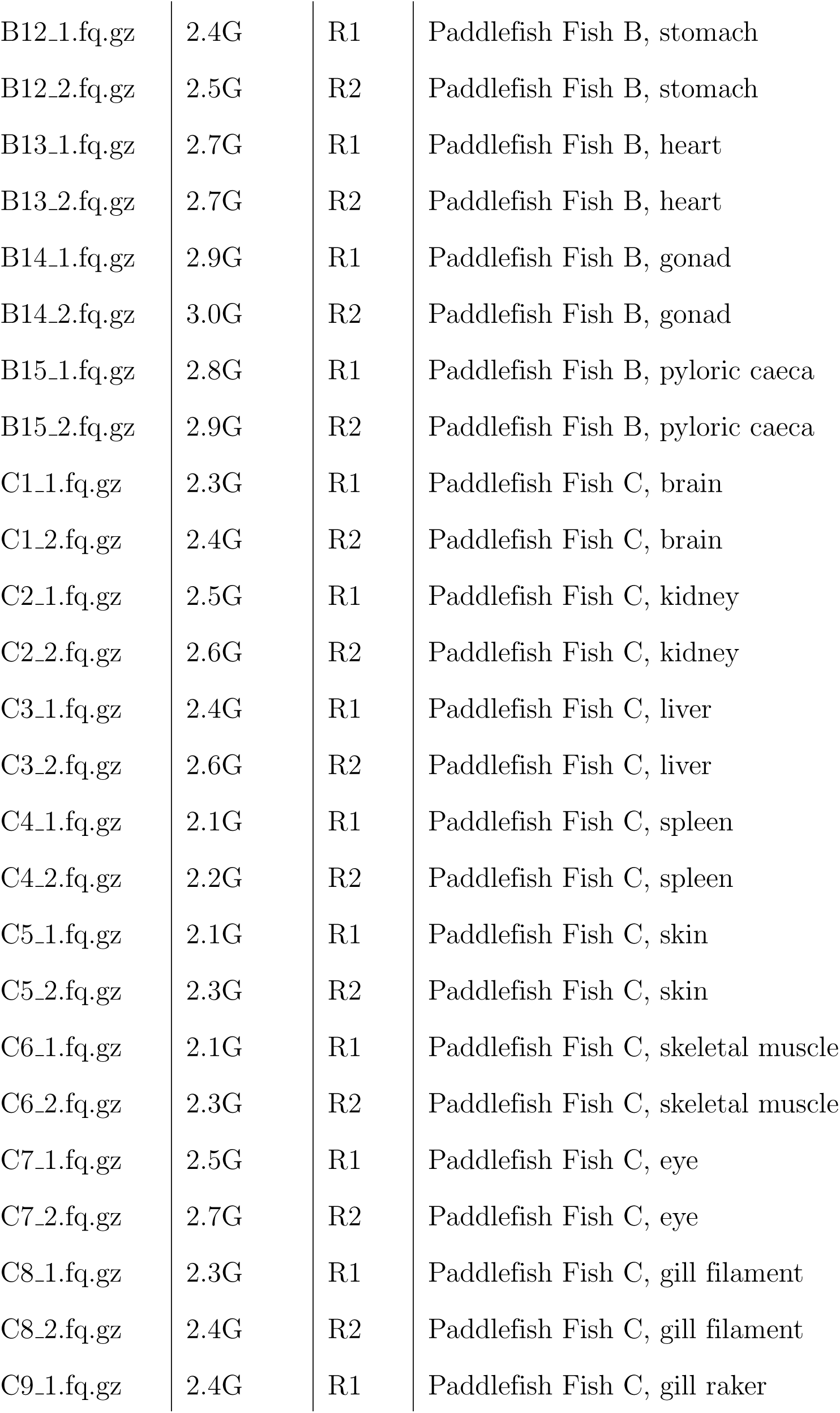
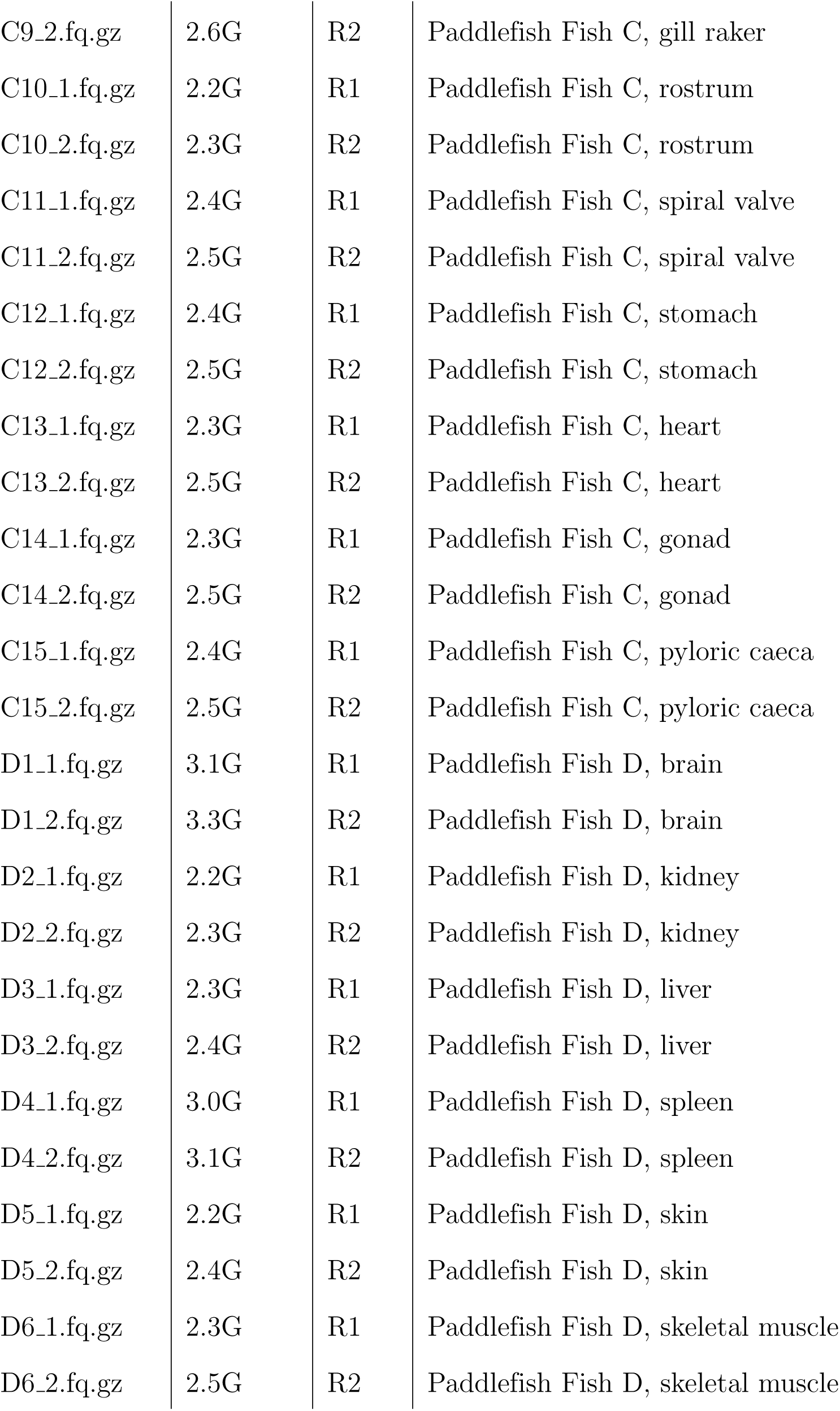
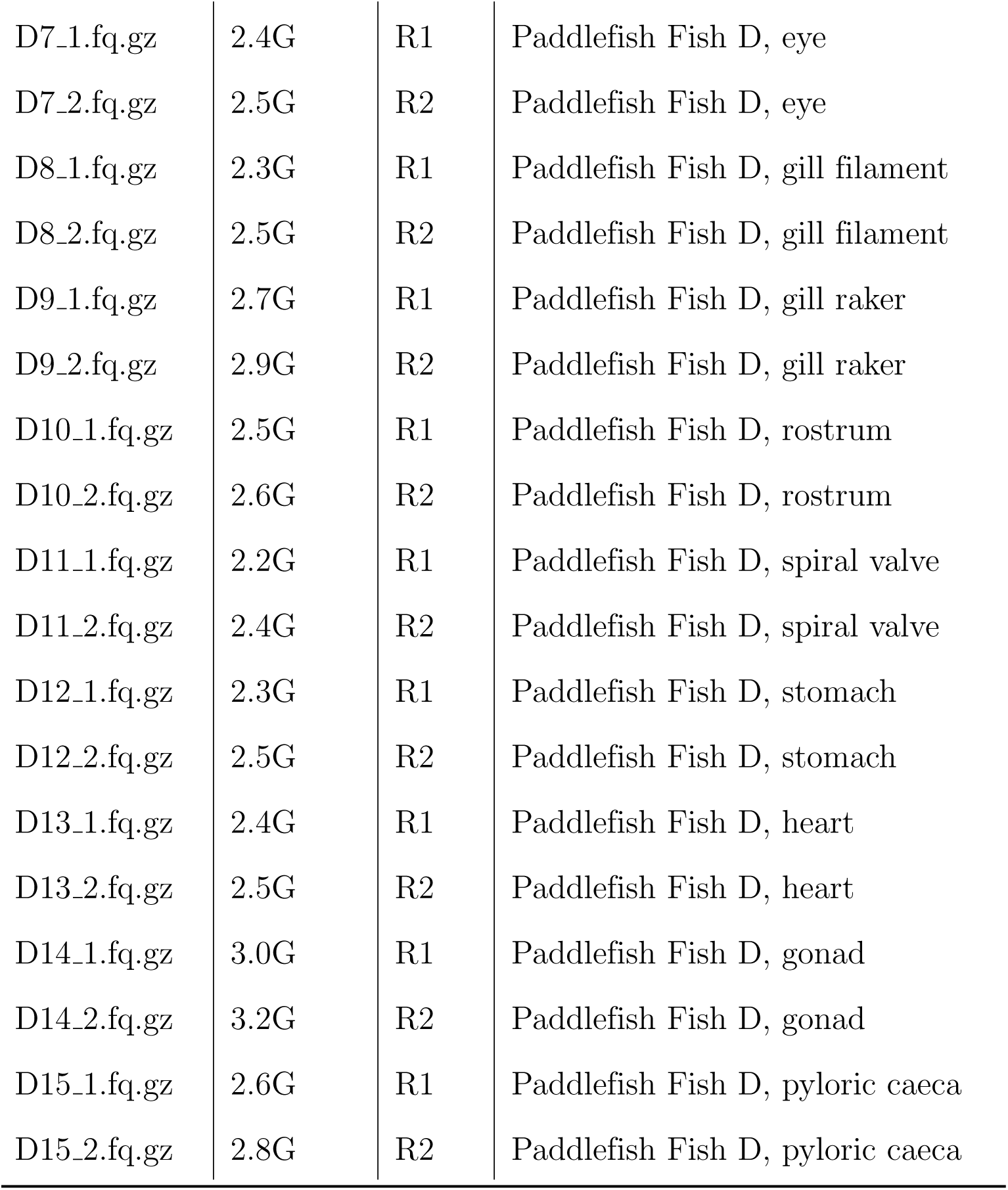
Summary of paired-end RNA-seq FASTQ files generated from *Polyodon spathula* (paddlefish) tissues deposited in the NCBI Sequence Read Archive (SRA) under BioProject accession PRJNA1512476. The dataset comprises 120 FASTQ files (60 paired-end libraries) from four individuals (A–D) and 15 tissue types per individual, with a total compressed size of approximately 296 GB.

**Figure S1.**
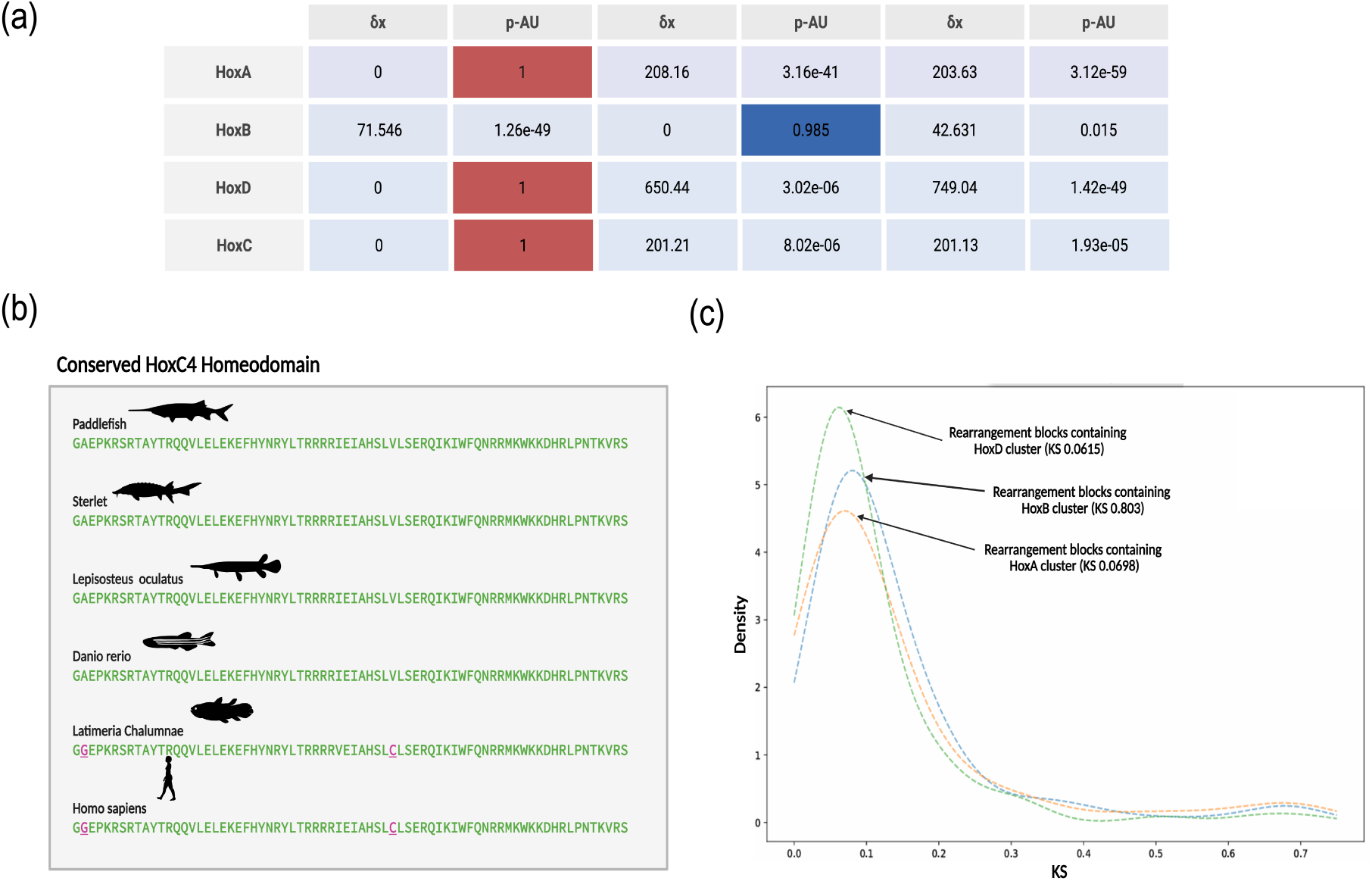
AU-test results for the concatenate Hox clusters in the sterlet and paddlefish (A) AU-test results for the concatenated gene trees (B) Alignment of itHoxC4 genes accross vertebrates showing clear conseravtion in the Paddlefish sequence (C) *K_s_* distributions for rearrangement blocks containing duplicated Hox clusters;*HoxA*, *HoxB*, and *HoxD* clusters in Sterlet.

**Table S5.** Hi-C QC. Summary of Hi-C sequencing, mapping, and library quality statistics for the paddlefish Hi-C dataset used for genome-wide TAD identification.

| Metric | Value |
| --- | --- |
| Sequenced reads | 298 M |
| Valid Hi-C contacts | 161 M |
| Interchromosomal | 68.8 M |
| Intrachromosomal $\geq 20$ kb | 19.7 M |
| Duplicates | 31 k |
| Self-ligation/dangling ends | Low |

**Figure S2.**
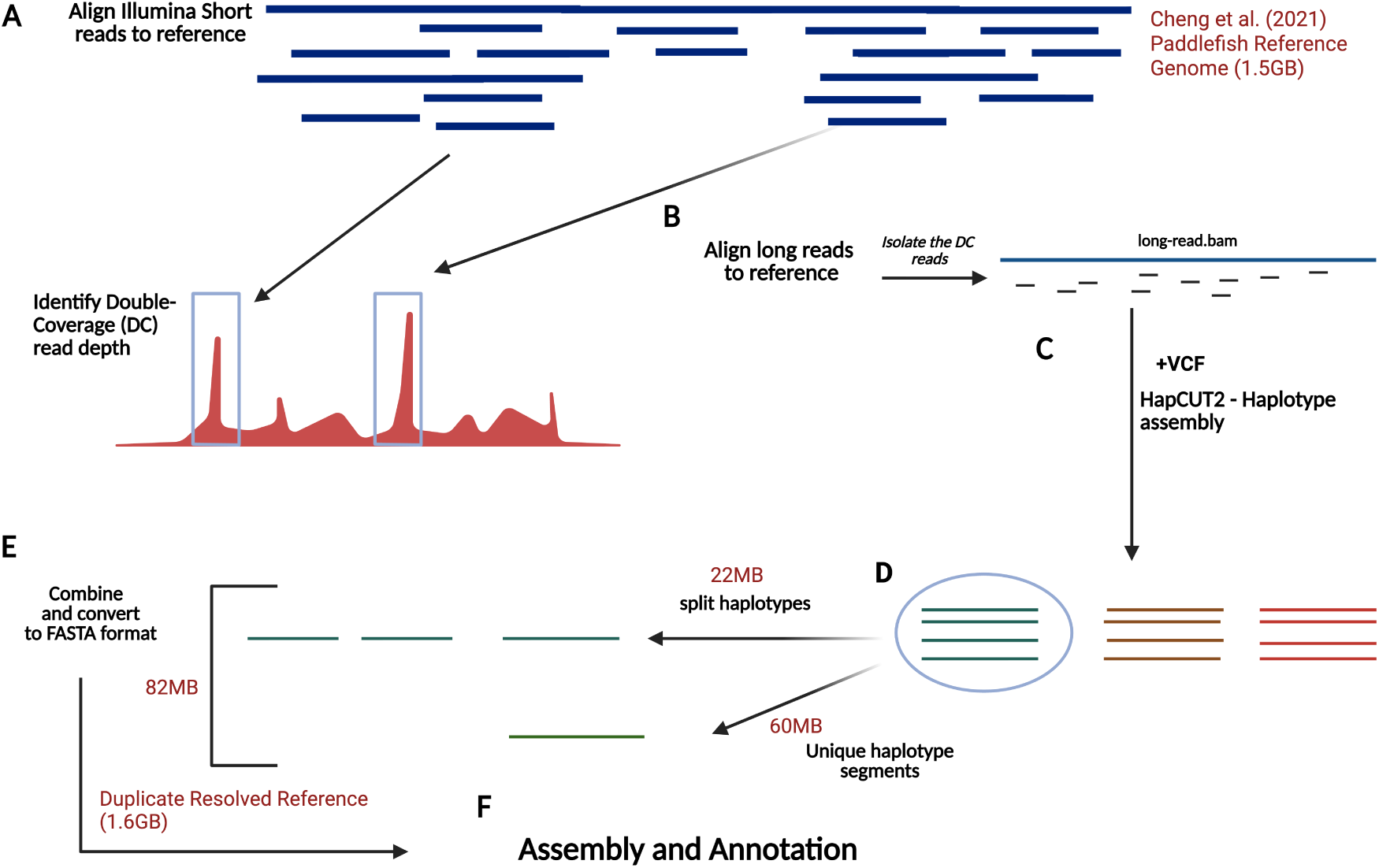
Schematic of the methodology used to resolve collapsed reads in the Cheng et al (2021), paddleFish Bssembly. (a) Illumina short reads are aligned to the reference, and regions with double coverage are identified using Mosdepth (v0.3.3) (66). (b) Double coverage (DC) regions are isolated from the mapped long reads (c) In conjunction with the DC long-read BAM file, a VCF generated with freebayes (26) is utilized as input for HapCUT2 ((20)) to reconstruct individual haplotypes within the double-coverage mapped long reads (d) For each double-coverage region, a haplotyped VCF file is generated. Some of these files contain more than one haplotypic segment These are our putative collapsed duplicates and so are split based on the haplotypic information from the VCF file (e) The split and unique regions are then integrated with the original genome fasta file (f) Subsequently, the duplicate resolved reference is scaffolded and annotated

